# Prolonged systemic inflammation worsens impairments to astrocyte Ca^2+^ and functional hyperemia in Alzheimer’s disease

**DOI:** 10.1101/2025.08.31.673380

**Authors:** Chang Liu, Kimia Sakha, Jaime Anton, Alfredo Cardenas-Rivera, Mohammad A. Yaseen

## Abstract

Chronic neuroinflammation in Alzheimer’s disease (AD) activates astrocytes—key regulators of both brain immunity and neurovascular coupling. The primed immune environment in AD brain also renders it highly susceptible to secondary systemic inflammatory challenges. Inflammatory activation drives phenotypic shifts in astrocytes that may compromise their ability to regulate cerebral blood flow. The capacity for inflammation-activated astrocytes to retain this regulatory function, however, remains unknown. To investigate astrocyte regulation of cerebral blood flow in AD brain and under systemic inflammation, we investigated astrocytic Ca^2+^ dynamics and functional hyperemia at rest and during brief and prolonged sensory stimulation in 12-month-old female APP/PS1dE9 mice. We further examined how a secondary systemic inflammatory challenge induced by low-dose, repeated injection of LPS modulates astrocytic signaling and neurovascular function. AD mice exhibited elevated spontaneous but diminished stimulation-evoked astrocytic Ca^2+^ activity, accompanied by impaired sustained functional hyperemia, particularly within the capillary network. LPS-induced systemic inflammation further suppressed both spontaneous and evoked astrocytic Ca^2+^ responses and attenuated functional hyperemia. Together, these findings reveal that inflammation disrupts astrocyte-dependent regulation of sustained neurovascular responses in the AD brain.

**HIGHLIGHTS:**

- Astrocytes in AD mice exhibit increased spontaneous Ca^2+^ signaling but cannot sustain stimulus-evoked Ca^2+^ release.
- Reduced astrocyte Ca^2+^ release during 30s functional brain activation correlates with impaired neurovascular coupling in both penetrating arterioles and capillaries of AD mice
- A secondary, 14-day systemic inflammatory challenge further suppressed functional hyperemia of 30 s stimulus–evoked astrocytic Ca^2+^ release in AD mice.
- A secondary inflammatory insult lasting 14 days reduced amyloid deposition in the AD brain.

## 1. INTRODUCTION

Vascular dysfunction and neuroinflammation are two major pathological features of preclinical Alzheimer’s disease (AD) and other diseases of the central nervous system (CNS) [1–4]. These notable AD hallmarks are coupled through their shared cellular components, with astrocytes performing especially critical roles in regulating both microvascular blood flow and brain immunity [5,6]. Astrocyte excitability is mediated primarily by Ca^2+^ signaling and is crucial for numerous immune, synaptic, metabolic, and hemodynamic signaling pathways of the “neuro-glio-vascular unit” (NGVU) [7–9]. In response to stress, injury, or disease-related alterations, astrocytes undergo phenotypic and morphological changes to a reactive state. Reactive astrocytes demonstrate altered excitability, dysregulated Ca^2+^ signaling, altered gene expression, metabolic shifts, and active release pro- or anti-inflammatory cytokines, and they contribute notably to AD progression and other CNS disorders [7,10,11]. In AD and other disorders, reactive astrocytes reportedly facilitate amyloid beta (Aβ) pathology and neural network dysfunction [7,8]. The impact of inflammatory astrocyte phenotype alterations on their contributions to cerebral blood flow regulation is presently unclear.

Neuroinflammation reportedly potentiates Aβ accumulation, cerebrovascular pathology, mitochondrial dysfunction, reactive oxygen species generation, and synaptic loss. Thus, it is increasingly recognized as a driving force and critical accelerator for the aggressive cycle of pathological deteriorations in preclinical AD and other neurodegenerative disorders [14,15]. Many investigators assert a nuanced influence of neuroinflammation, citing evidence of both harmful and beneficial effects of an inflammatory stimulus during disease progression [16,17]. Several studies also highlight the importance of the brain’s initial inflammatory state on subsequent outcomes to a secondary inflammatory threat. At early stages of AD or other neurodegenerative diseases, acute periods of illness or systemic inflammation can catalyze chronic neuroinflammation and exacerbate neurotoxicity and functional decline [18–21]. Collectively, these views motivate investigations to explore the effects of enhancing or repressing a pro-inflammatory response on chronic cerebral pathologies [22].

Previous studies also show that acute systemic inflammation disrupts cerebral hemodynamics in healthy rodent brains and human patients with small vessel disease and traumatic brain injury [23–25]. To precisely elucidate the intricate relationship between neuroinflammation and vascular dysfunction in preclinical AD, a detailed understanding of neuroinflammation’s impact on astrocyte Ca^2+^-mediated microvascular hemodynamics is essential. In this study, we investigated the effects of amyloid-induced inflammation coupled with a prolonged systemic inflammatory threat on astrocyte Ca^2+^ signaling and cerebral blood flow. 12-13 month-old female APPswe/PS1dE9 mice and their wildtype littermates (n=8 each) were exposed to a 14-day period of systemic inflammation induced by intraperitoneal injection of the bacteria-derived endotoxin lipopolysaccharide injection (LPS) [26]. Using intravital two-photon microscopy, we longitudinally measured Ca^2+^ release from astrocytes and microvascular hemodynamics in penetrating arterioles and capillary branches under resting state conditions and during functional brain activation in the awake mouse brain. Our observations demonstrate that Aβ-induced inflammatory disruptions notably alter astrocyte Ca^2+^ and sensory-evoked microvascular responses. A prolonged period of LPS-induced inflammation further exacerbated astrocyte Ca^2+^ signaling and the hyperemic response to longer, 30 second functional stimulation. Immunofluorescence imaging showed reduced levels of cortical and hippocampal Aβ after 14-days of induced inflammation. The results delineate the interdependent relationships between astrocyte Ca^2+^ signaling, neuroimmune and neurovascular dynamics, and Aβ pathology during preclinical AD’s multifactorial progression.

## 2. RESULTS

Among its many functions, astrocyte Ca^2+^ is an important modulator of basal blood flow and functional hyperemia in the brain [6,27]. To determine whether Aβ-induced inflammatory astrogliosis alters astrocyte Ca^2+^-mediated contributions to cerebral blood flow regulation, we measured astrocyte Ca^2+^ dynamics and microvascular dilation under resting-state conditions and during sensory-evoked functional hyperemia in the awake mouse brain at age 12-13 months in APPswe/PS1dE9 and wildtype mice. We then explored if a 14-day LPS-induced neuroinflammation challenge alters astrocyte calcium signaling and its contributions to neurovascular coupling. Systemic injection of the bacteria-derived endotoxin lipopolysaccharide (LPS) has been extensively used as an effective model to study neuroinflammation in neurodegenerative disease [26].

Recent studies demonstrate that astrocyte Ca^2+^ contributes significantly to sustaining functional hyperemia in response to prolonged, but not brief, periods of cortical stimulation [28]. Motivated by these findings, we used two-photon imaging to measure brain hemodynamics and astrocyte Ca^2+^ release in response to both brief (3s) and sustained (30s) whisker stimulation.

Intravital two-photon microscopy measurements were performed in cortical layers I and IV of awake mice. Data from both layers were combined.

### 2.1 Pre-inflammation: Hyperactive astrocytes and reduced stimulus-evoked astrocyte Ca^2+^ in AD mice

We first evaluated resting-state spontaneous cortical astrocyte Ca^2+^ by imaging GCaMP6f-labelled astrocytes in awake mice. Raster scanning was performed in ∼88 μm^2^ cortical regions surrounding penetrating arterioles for 10 min at ∼ 4.4 Hz (Fig. 1A). Dynamic Ca^2+^ transients were quantified and analyzed from the soma, processes, and endfoot compartments of astrocytes. In AD mice, we observed ∼15 – 40% higher frequencies of Ca^2+^ spikes compared to age-matched wildtype littermates, with significantly higher Ca^2+^ frequency in the soma and process compartments (Fig. 1B-C, left, p-val = 0.03 soma, p-val = 0.048 processes). No differences in the Ca^2+^ amplitudes were observed between AD and WT mice. Compared to WT littermates, the mean duration of the astrocytic Ca^2+^ spikes in AD mice was significantly lower (26%, p-val < 1e-4) in astrocyte processes. Shorter Ca^2+^ durations were also observed in the soma and endfoot compartments in AD mice; however, the differences did not reach statistical significance (Fig. 1C).

**Fig. 1.**
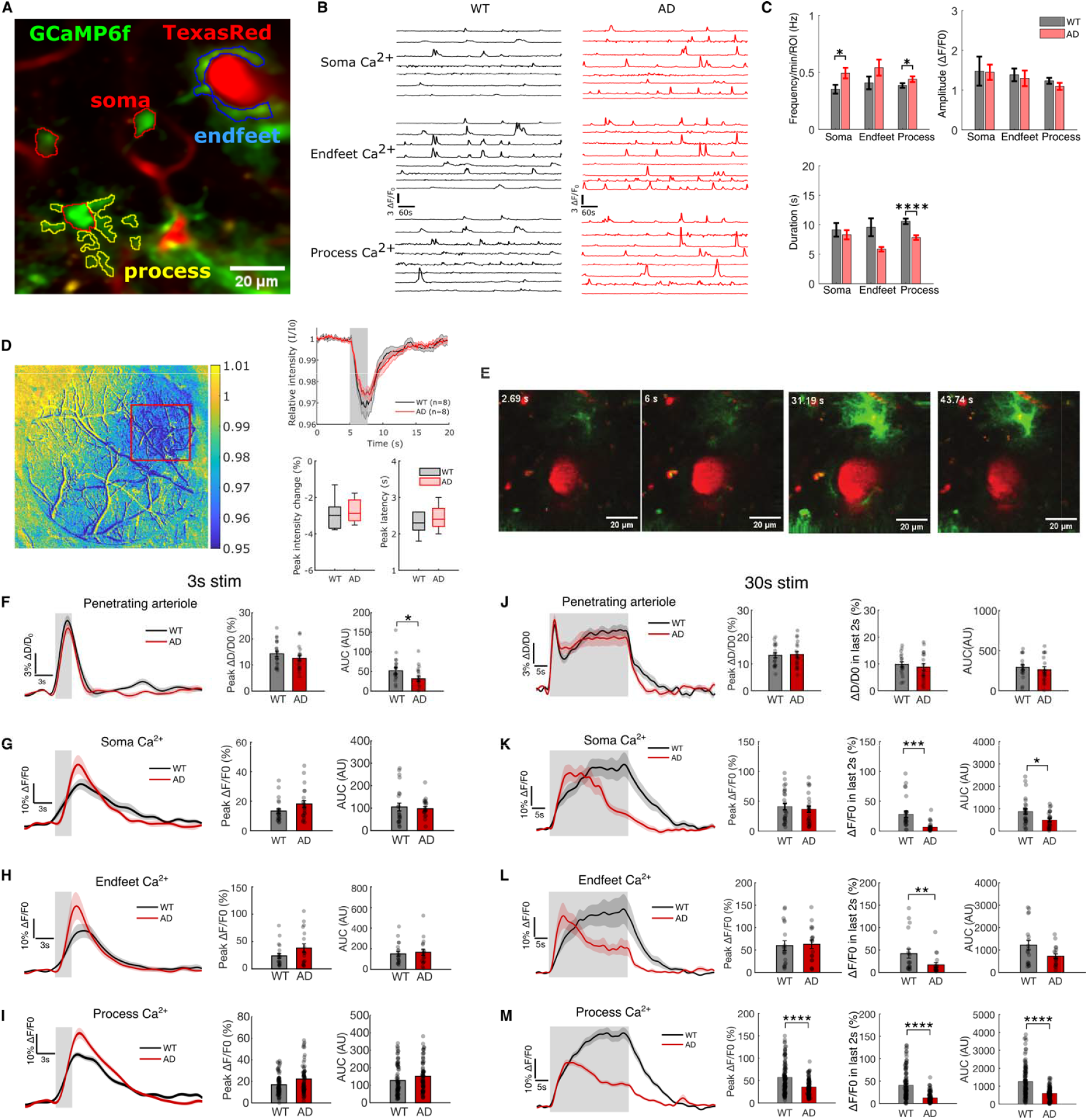
Spontaneous astrocyte Ca^2+^ and sustained sensory stimulation in AD and WT mice barrel cortex. **(A)** 2D, raster scan image (average intensity projection of a 10-minute time series) showing vasculature labelled with Texas-Red and astrocyte microdomains expressing GCaMP6f. Annotated regions are soma (red), endfeet (blue), and process (yellow). **(B)** Representative measurements of spontaneous Ca^2+^ release in cortical astrocytes cell compartments recorded for 10 mins. **(C)** Amplitude, Frequency and Duration of Ca^2+^ spikes at different astrocyte ROIs. For WT mice, 25 somas and 156 processes were measured from 8 mice, and 14 endfoot were measured from 7 mice. For AD mice, 30 somas, 101 processes, and 12 endfoot were measured from 8 mice. **(D)** OISI measurement of cerebral hemodynamics in response to 3s whisker stimulation in left barrel cortex in WT and AD mice. **(E)** Representative two-photon raster scan images of diving arteriole dilation and astrocyte Ca^2+^ release during a 30s whisker stimulation trial. **(F-I):** Trial-averaged profiles of arteriole dilation and astrocyte Ca^2+^ release in response to 3s whisker stimulation and quantification of the peak response. **(J-M):** Trial-averaged profiles of arteriole dilation and astrocyte Ca^2+^ release in response to 30s whisker stimulation and the quantification of the peak response. Difference between WT and AD were tested using rank-sum test.

Next, we investigated whether amyloid pathology impacts functional hyperemia. We first applied optical intrinsic signal imaging (OISI) during a 3 s whisker stimulus to locate the activation center (Fig. 1D). We measured astrocyte Ca^2+^ dynamics and diameter of penetrating arterioles while stimulating the mouse barrel cortex via pneumatic whisker deflection. Our OISI measurement revealed slight, but not significant, differences in peak cerebral blood volume response (< −1%) in AD mice relative to WT (Fig. 1D). The activated region was then imaged via TPM using both raster scanning (4.4 Hz, Fig 1E) and rapid line scanning (∼360 – 100 Hz, downsampled to 20 Hz during data processing).

In response to a short ∼3s functional stimulus, AD and WT mice demonstrated different responses for arterial dilation and Ca^2+^ signaling within astrocyte compartments. The area under the curve (AUC) for arterial dilation response was significantly lower in AD mice (40% difference, p = 0.029), while arterial peak dilation, rise times, and latency were similar between cohorts (Fig.1F and supplementary Fig. 1). Astrocyte Ca^2+^ kinetics varied in different astrocytic microdomains, as reported previously [29]. Peak amplitudes of stimulus-induced Ca^2+^ release in AD mice appeared higher in all astrocyte compartments compared to WT mice; however, the differences did not achieve statistical significance (Fig. 1 G-I, soma p = 0.23, endfeet p = 0.21, processes p= 0.054). AD mice demonstrated significantly shorter Ca^2+^ responses in astrocytic endfeet (supplementary Fig. 1, p = 7e-4). In both cohorts, we consistently observed that the short 3s whisker stimulus evoked slower astrocyte Ca^2+^ responses relative to the arteriole response. Arterioles dilated to their maximum approximately 2.1 seconds after the stimulus onset, while all astrocyte Ca^2+^ release required 4 - 4.5 s to reach their peak values (∼ 1-1.5 s after stimulus cessation). Our results indicate that astrocytes in the AD mice brain respond slightly faster and more pronouncedly than WT mice to brief sensory stimulation.

A longer 30s whisker stimulus yielded more striking differences between AD and WT mice, particularly in astrocyte Ca^2+^ dynamics. Response profiles exhibited biphasic features for both arterial dilation and astrocyte Ca^2+^ release - including a rapid rising phase followed by a prolonged plateau phase. Consequently, we analyzed dilation and Ca^2+^ release both over the full stimulus duration (all 30 seconds) as well as the late phase (the final 2 seconds). Again, AD and WT mice experienced comparable peak diameter in arterioles within the first 3 seconds of stimulation. Following peak dilation, arterioles constricted slightly and plateaued to a smaller diameter (Fig. 1J). For both phases, we observed no significant differences in arterial dilation rise times, maximal diameter, or AUC between AD and WT mice. However, following stimulus cessation, we observed significantly faster constriction to baseline diameter in AD mice (Supplementary Fig. 1E, Δt = 5.04s, p = 0.03). Astrocyte Ca^2+^ responses differed more substantially between cohorts for the 30 s stimulus (Fig. 1 K-M & Supplementary Fig.1 F-H). In astrocyte soma and endfeet, both AD and WT mice achieved similar peak values of Ca^2+^ release, but AD mice achieved their peak significantly earlier. In WT mice, astrocytes persistently released Ca^2+^ from all compartments until the stimulus cessation, while Ca^2+^ release in AD astrocyte peaked rapidly during the first ∼5s of stimulation and diminished markedly thereafter. Within processes, AD mice demonstrated significant lower peak Ca^2+^ release (Fig. 1M & Supplementary Fig. 1H, p < 1e-4). For all astrocyte compartments, Ca^2+^ release was significantly lower in AD mice during the final 2s of the stimulus (Fig.1 K-M, soma p = 1e-4, endfeet p = 7e-4, processes p < 1e-4). Our observations indicate a compartmentalized influence of AD pathology to astrocyte Ca^2+^. In all microdomains, AD mice could not sustain astrocytic Ca^2+^ release in response to longer 30 s sensory stimulation, indicating reduced Ca^2+^ storage.

The influence of astrocytes on microvascular blood flow reportedly extends beyond penetrating arterioles to laterally-oriented pre-capillary arterioles and capillaries [29,30]. In this dense network of small, more prevalent microvessels, even modest alterations to vessel diameter can substantially impact vascular resistance and total blood flow [31]. To examine how amyloid pathology and associated hyperactive astrocytes affect hemodynamics in these vessels, we performed rapid two-photon line scan measurements to measure capillary diameter and red blood cell speed. Vessel dilation and RBC speed in laterally-oriented arteriole branches and capillary branches extending from the downward penetrating arteriole (1^st^ order to 4^th^ order) were measured and downsampled to a rate of 20Hz. The vessels were further grouped as precapillary arteriole (1^st^ to 2^nd^ order) and high-order capillaries (≥3^rd^ order) (Supplementary Fig. 2A). Baseline vessel diameters were computed as the mean diameter between 1.5 to 4.5s of the 5s baseline period (Supplementary Fig. 2B). In AD mice, we observed significantly faster RBC speed in high-order capillaries compared to WT mice, suggesting higher blood flow and lower vascular resistance (Supplementary Fig. 2C). For the short 3s stimulus duration, we observed no differences in relative peak dilation for pre-capillary arterioles or capillaries (Supplementary Fig. 2D-F). For a longer 30 second stimulus, however, responses from higher-order capillaries in AD mice had significantly lower diameters during the last 2 seconds and AUC (Supplementary Fig. 2 H).

### 2.2 LPS-induced inflammation reduces spontaneous astrocyte Ca^2+^ and impairs neurovascular coupling

To evaluate how prolonged systemic inflammation alters astrocyte’s Ca^2+^ signaling and its contributions to CBF regulation, animals underwent a 14-day LPS-induced inflammation challenge. We used 2-photon imaging to measure astrocyte Ca^2+^ dynamics and microvascular dilation under resting-state conditions and during functional hyperemia in awake AD and WT mice. The same imaging fields of view were imaged on days 0, 7, and 14 of LPS administration.

Fig. 2A displays example measurements of resting-state, spontaneous Ca^2+^ release from astrocytic compartments over the course of the 14-day inflammatory threat. Once again, raster scanning was performed in ∼88 μm^2^ cortical regions surrounding penetrating arterioles for 10 min at ∼ 4.4 Hz. We observed no discernible differences in astrocyte morphology from days 0 – 14. Dynamic Ca^2+^ transients were quantified and analyzed from the soma, processes, and endfeet compartments of astrocytes (Fig. 2 B & C). During the 14-day period, we observed variations in spontaneous Ca^2+^ fluctuations in astrocytes in both cohorts, with some cell compartments demonstrating no Ca^2+^ transients over the 10-minute measurement. Fig. 2D illustrates how the proportion of these ‘silent,’ quiescent cell compartments changed during the inflammatory period. Among the cells that remained active in both AD and WT cohorts, the frequency of spontaneous Ca^2+^ spikes progressively lowered during the 14-day period within astrocyte soma and process compartments (Fig. 2E&G). No substantial changes were observed in endfeet Ca^2+^ (Fig. 2F).

**Fig. 2.**
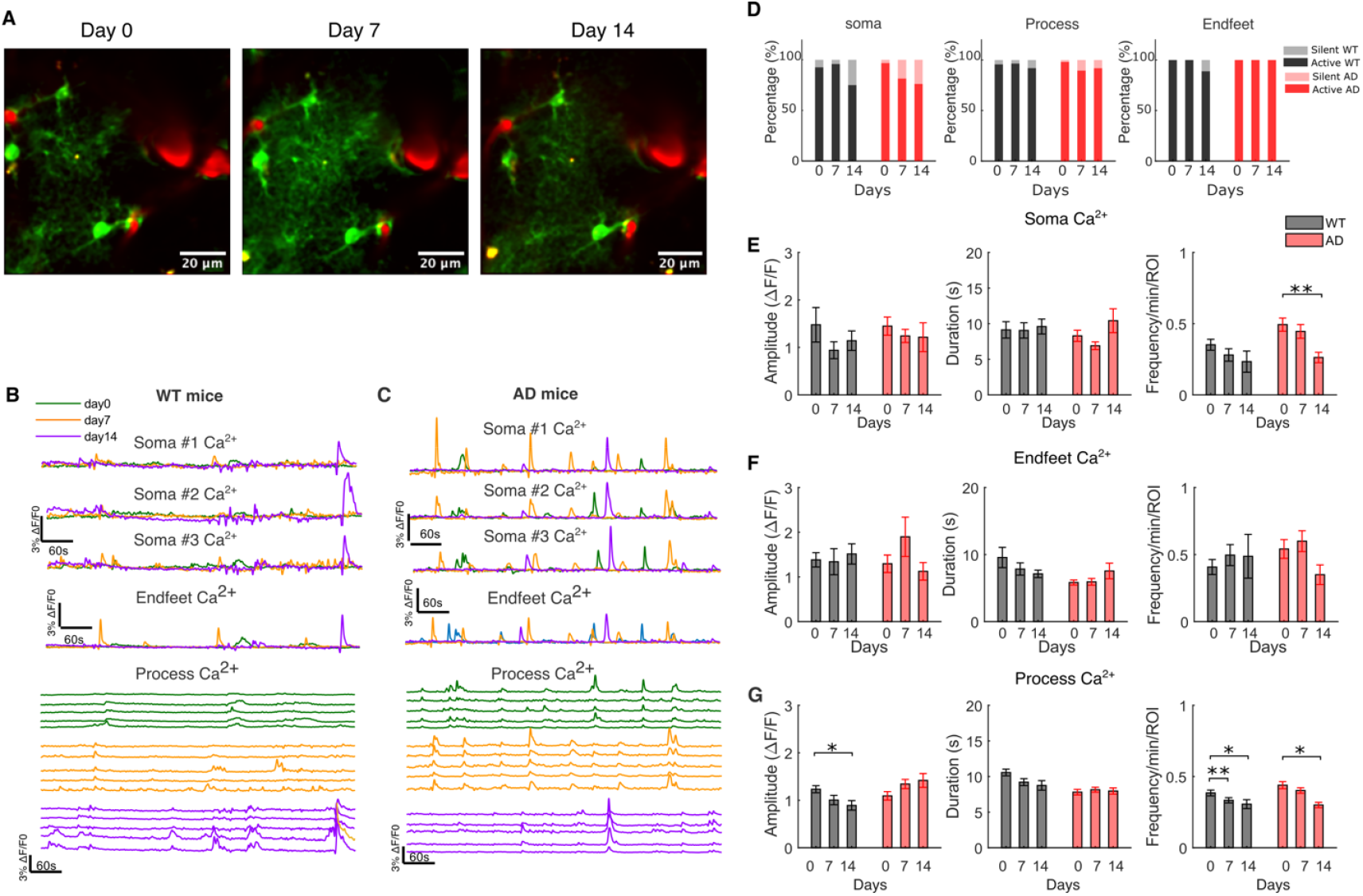
LPS induced inflammation decreased spontaneous Ca^2+^ release in astrocytes in AD mice cortex. **(A)** Raster scan image of cortical vasculature and astrocytes at different days during 14-days of LPS-induced systemic inflammation (average intensity projections from 10-minute time series). **(B)** Spontaneous Ca^2+^ release form astrocyte compartments in WT and AD mice cortex measured on pre-LPS injection (day 0), after 7 and 14 days of LPS injection (day 7 and day14). **(D)** Variations in spontaneous Ca^2+^ activity from astrocyte compartments during the 14-day period. **(E-G)** Quantification of astrocyte Ca2+ spikes under the impact of LPS induced inflammation. Linear mixed effect model was used to test the effect of LPS injection. WT mice day0 measurement: 25 somas, 156 processes and 14 endfoot measured from 8, 8, and 7 mice, respectively. WT mice day7 measurement: 23 somas, 111 processes and 17 endfoot measured from 8, 8, and 8 mice. WT mice day14 measurement: 12 somas, 46 processes and 8 endfoot measured from 4, 6 and 6 mice. AD mice day0 measurement: 30 somas, 101 processes and 12 endfoot measured from 8, 8, and 8 mice. AD mice day7 measurement: 22 somas, 103 processes and 11 endfoot measured from 7, 8 and 7 mice. AD mice day 14 measurement: 16 somas, 68 processes and 8 endfoot measured from 5, 7 and 5 mice.

To determine whether prolonged systemic inflammation impacts astrocytic contributions to functional hyperemia, we again examined dilation of penetrating arterioles and astrocyte Ca^2+^ release in response to brief (3s) and sustained (30 s) pneumatic whisker stimulation. For short 3s whisker stimulation, arteriole dilation responses did not change appreciably during the 14-day period for both AD and WT mice (Fig. 3A), while astrocyte Ca^2+^ responses varied significantly. In the astrocyte soma of WT mice, Ca^2+^ release parameters showed reductions in peak response and AUC between days 0 and 7 (Fig. 3B, p-val = 0.036 and p-val = 0.047, respectively). For both AD and WT mice, systemic inflammation substantially altered Ca^2+^ release in astrocyte processes, provoking significant reductions in peak Ca^2+^ release, AUC, and rise time (Fig. 3D). Interestingly, the inflammatory threat induced opposing effects in AD and WT mice within astrocyte endfeet. In WT mice, we observed significant increases in peak Ca^2+^ of endfeet and shorter response duration. Conversely, in AD mice, endfeet showed reduced peak Ca^2+^ and significantly longer rise times, but longer duration (Fig. 3C & Supplementary Fig. 3).

**Fig. 3.**
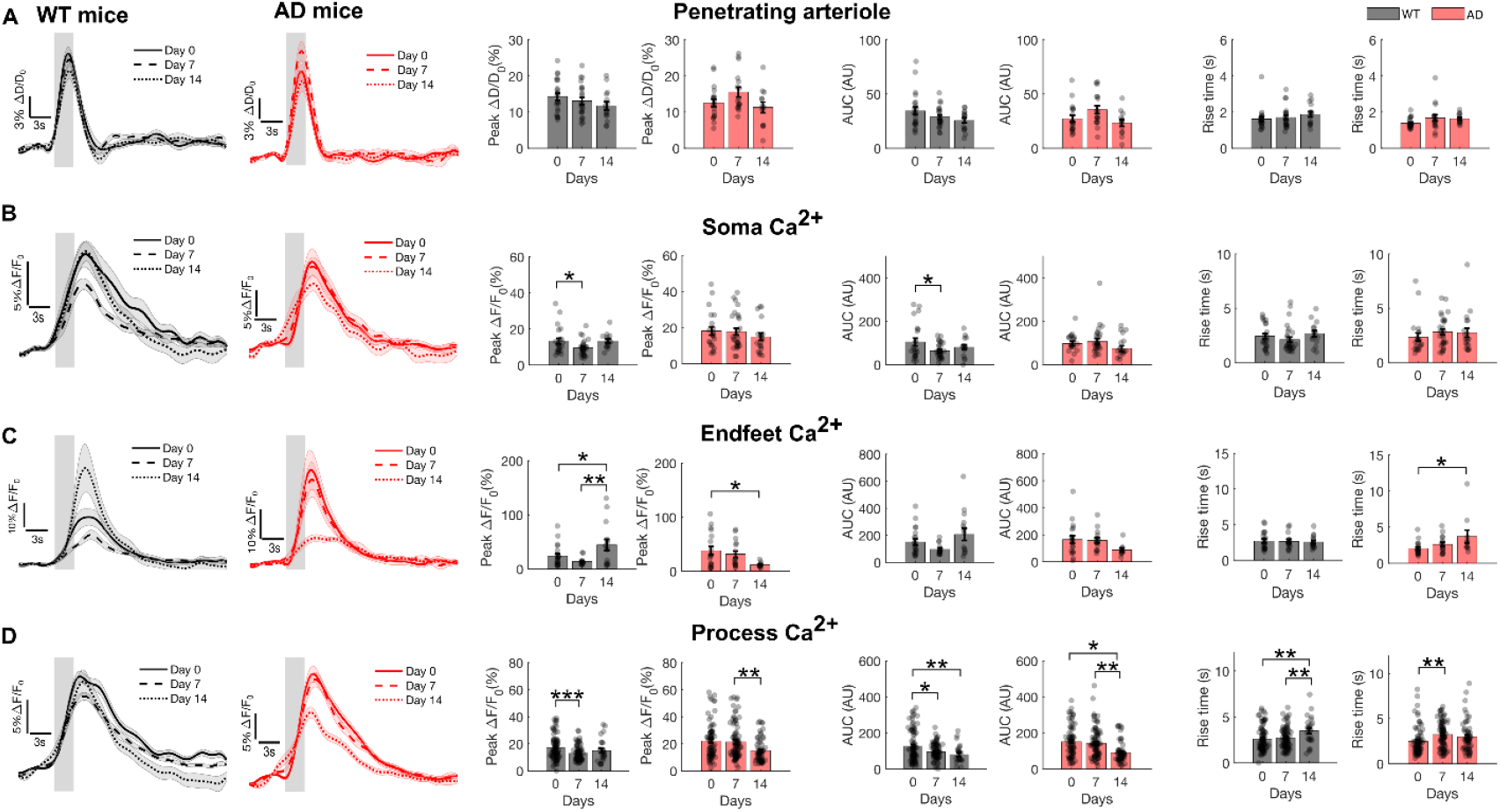
Effect of LPS induced inflammation on penetrating arterial dilation and astrocyte Ca^2+^ release in response to brief whisker 3s stimulation. (**A)** Trial-averaged profiles and calculated features of stimulus-induced penetrating arteriole dilation in WT and AD mice on day 0 (WT: N=22 arterioles, n=7 mice; AD: N=19 arterioles, n=8 mice), day7 (WT: N=22 arterioles, n=8 mice; AD: N=17 arterioles, n=8 mice), and day14 of LPS injection (WT: N=15 arterioles, n=5 mice. AD: N=13 arterioles, n=6 mice). (**B-D)** Trial-averaged profiles and calculated features of stimulus-induced, compartment-specific astrocyte Ca^2+^ release in WT and AD mice on day 0 (WT: N=23 soma from 8 mice, N=20 endfoot from n=7 mice, N=88 processes from n= 8 mice. AD: N=21 soma from 7 mice, N=18 endfoot from 8 mice, N=77 processes from 7 mice), day7 (WT: N=29 soma from 7 mice, N=15 endfoot from n=6 mice, N=86 processes from 8 mice. AD: N=28 soma from 7 mice, N=16 endfoot from 8 mice, N=86 processes from 8 mice) and day14 of LPS injection (WT: N=15 soma from 5 mice, N=15 endfoot from n=5 mice, N=25 processes from 5 mice. AD: N=19 soma from 5 mice, N=10 endfoot from 6 mice, N=62 processes from 6 mice).

For longer 30 second whisker stimulation, the 14-day inflammatory threat provoked more pronounced alterations to WT mice than AD mice. Again, more significant inflammation-induced alterations were observed in astrocytes than arterioles. In both AD and WT mice, arteriole dilation was reduced during the later plateau phase at days 7 and 14 (Fig. 4A) and AD mice showed significantly slower restoration to baseline diameter (supplementary Fig. 4A). In WT mice, peak Ca^2+^ release and rise times during the entire 30 s stimulation all significantly decreased in soma and processes over the 14-day inflammatory period, and peak latency was also significantly decreased in processes. AUC significantly lowered within all compartments (Fig. 4 B-D). Maximum Ca^2+^ release during the late phase of stimulation were also significantly reduced in all compartments (Fig. 4 B-D). In AD mice, the inflammatory threat provoked significant reductions in peak Ca^2+^ release from all astrocyte compartments. The AUC for responses in astrocyte endfeet also decreased significantly during the inflammation period in AD mice, while astrocyte processes showed significant reductions in AUC, peak latency, duration, and the average Ca^2+^ release during the later plateau phase (Fig. 4 D & supplementary Fig. 4D). Overall, our observations at the late phase of stimulation show reduced endfeet Ca^2+^ release that correlate well with mildly reduced penetrating arteriole dilation and slower post-stimulus constriction. The results align well with prior studies demonstrating that astrocyte Ca^2+^ does not initiate arteriole dilation but is required to sustain dilation and mediate recovery from prolonged functional stimulation [28].

**Fig. 4.**
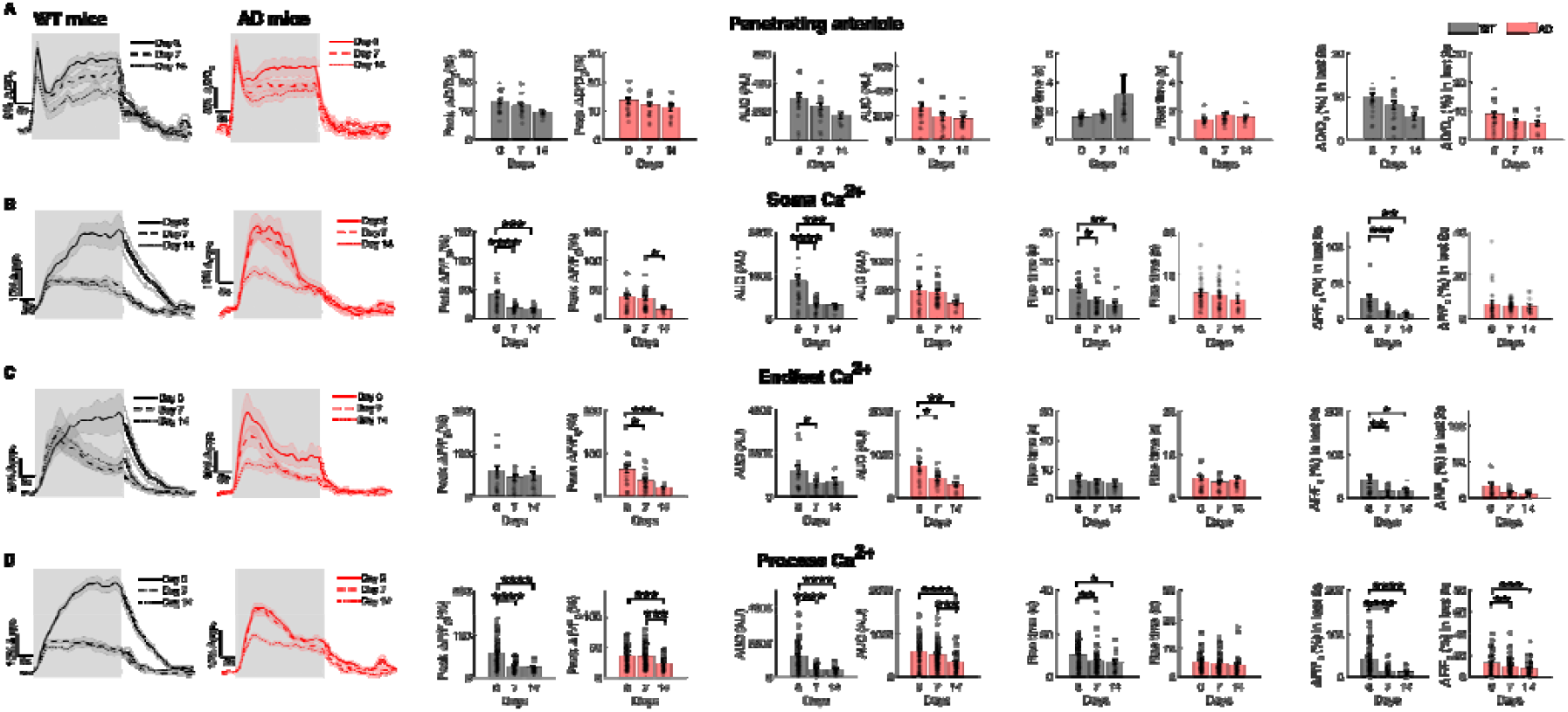
Effect of LPS induced inflammation on penetrating arteriole dilation and astrocyte Ca^2+^ release in response to 30 second whisker stimulation. (**A)** Trial-averaged profiles and calculated features of stimulus-induced penetrating arteriole dilation in WT and AD mice on day 0 (WT: N= 18 arterioles, n=7 mice. AD: N=18 arterioles, n=8 mice), day7 (WT: N=18 arterioles, n=8 mice. AD: N=19 arterioles, n=8 mice), and day14 of LPS injection (WT: N= 9 arterioles, n=5 mice, AD: N=14 arterioles, n=6 mice). (**B-D)** Trial-averaged profiles and calculated features of stimulus-induced, compartment-specific astrocyte Ca^2+^ release in WT and AD mice on day 0 (WT: N=26 soma from 8 mice, N=19 endfoot from n= 7 mice, N= 114 processes, from n=8 mice. AD: N=24 soma from 7 mice, N=18 endfoot from 8 mice, N=94 processes from 7 mice), day7 (WT: N=28 soma from 8 mice, N=18 endfoot from n=8 mice, N=98 processes from 8 mice. AD: N=32 soma from 8 mice, N=19 endfoot from 8 mice, N=114 processes from 8 mice) and day14 of LPS injection (WT: N= 14 soma from 5 mice, N=11 endfoot from n= 5 mice, N=24 processes from 5 mice. AD: N=12 soma from 5 mice, N=13 endfoot from 6 mice, N=71 processes from 6 mice).

We then examined how inflammation impacts stimulus-evoked dilation of laterally-oriented pre-capillary arterioles and capillaries. Using rapid line scan measurements, we measured kinetic profiles of stimulus induced vessel dilation and RBC velocity in lateral arterioles and capillary branches. The line scan measurements from enveloping astrocyte endfeet and neighboring processes further indicate that systemic inflammation impairs their stimulus-evoked Ca^2+^ release; however, we observed no significant LPS-induced alterations in vessel dilation or RBC velocity from capillaries or pre-capillary arterioles (supplementary figures 5-7).

### 2.3 Ex vivo immunofluorescence analysis of LPS-induced inflammation and A *β*_1-42_ accumulation

To assess neuroinflammation after LPS administration, we harvested brains from subgroups of mouse cohorts for immunofluorescence (IF) staining of GFAP, Iba-1, and Aβ_1-42_ (n = 3 per cohort). We also analyzed a sham group of mice that received 14 days of saline injections (n=3 for AD, n=2 for WT) and an additional group of mice that received 7 days of LPS injection (n=2 for AD, n=3 for WT). Note that the mice received 7 days of LPS injection were not used for *in vivo* imaging. Fig. 5A displays representative confocal images of immunofluorescence staining. We quantified GFAP expression in astrocytes and IBA1 expression in microglia by calculating the percentage area of GFAP and Iba-1 expressed cells in the confocal images (Fig. 5B). In the control group, AD mice showed significantly higher GFAP expression in both cortex and hippocampus compared to WT mice, indicating a pre-existing neuroinflammatory state in AD mice. IBA-1 expression is slightly, but not significantly, higher than in AD control compared to WT control mice. After 7 days of LPS injection, GFAP and IBA1 expression in the WT brain both increased, with a statistically significant increase observed in GFAP expression. Interestingly, GFAP expression in AD astrocytes decreased after 7 days of LPS injection while microglial expression of Iba-1 increased. Iba-1 expression reduced after 14 days in both cohorts. This trend of recovery agrees well with our previous study exploring cortical oxygen extraction during 14 days of LPS injection [32]. The results could reflect immune tolerance induced by the repeated injection of low-dose LPS [33–35]. Fig. 5C shows the representative widefield fluorescence imaging of Aβ_1-42_ in the cortex and hippocampus of AD mice. Strikingly, 14 days of LPS injection yielded reduced A*β*_1-42_ load in AD mice brain, showing statistical significance in cortex and a strong trend in the hippocampus (Fig. 5D). LPS injection induced A*β*_1-42_ reduction was also reported by other studies [36–38]. Higher A*β*_1-42_ load was observed in the cortex compared to hippocampus (Fig. 5D), agreeing well with previous reports in APPswe/PS1dE9 mice [39,40].

**Fig. 5.**
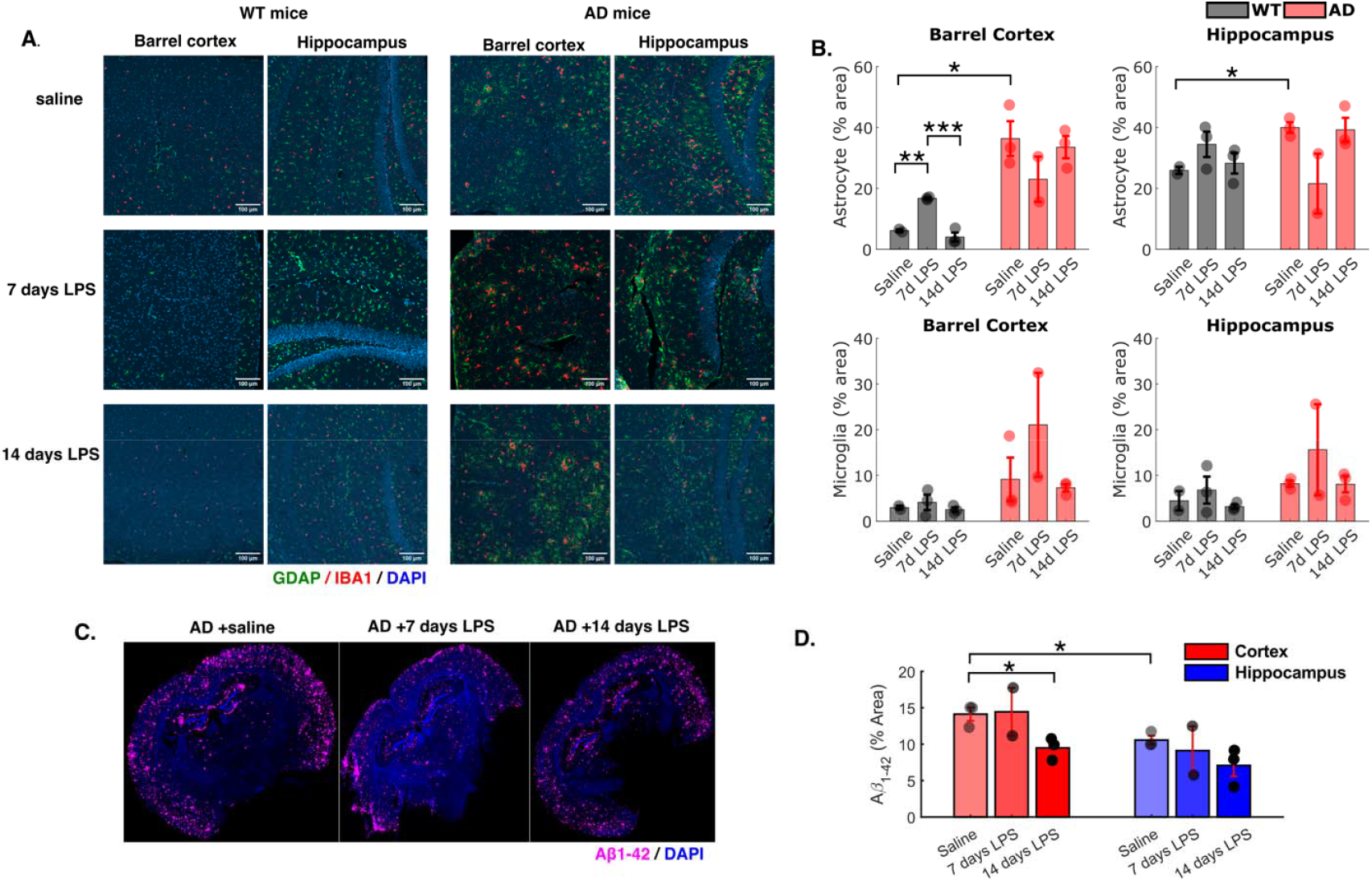
*Ex vivo* immunofluorescence imaging. **(A)** Representative confocal immunofluorescence image of GFAP and Iba-1expression in barrel cortex and hippocampus in WT and AD mice. Scale bar = 100 μm. **(B)** Quantification of GFAP and IBA-1 expression in barrel cortex and hippocampus. **(C)** Immunofluorescence image of in AD and WT mice that received 14 days of saline injection, 7 days of LPS injection and 14 days of LPS injection. **(D)** Quantification of in cerebral cortex and hippocampus. Multiple ROIs were quantified and averaged for each animal (percentage area, mean sem). One-way ANOVA with Tukey’s post hoc test was used in **(B)** and **(D)** to test the difference between saline, 7 days LPS and 14 days LPS injection. Differences between WT and AD saline control group in **(B)** and **(D)** were tested using Student’s t test. *p<0.05, **p<0.01, ***p<0.001.

## 3. DISCUSSION AND CONCLUSION

Our findings demonstrate that amyloid-induced astrocyte reactive alterations disrupt their Ca^2+^-mediated roles in modulating basal cerebral blood flow and stimulus-evoked functional hyperemia. In 12 to 13-month old female APPswe/PS1dE9 mice and their wildtype littermates, a prolonged 14-day period of LPS-induced systemic inflammation further diminishes Ca^2+^ release from astrocytic soma, processes, and endfeet, and it impairs stimulus-evoked dilation of penetrating arterioles, but not capillaries. Inflammatory disruptions to functional hyperemia are more pronounced for longer stimuli relative to shorter stimuli (30s vs 3s), reflecting the critical roles of astrocytes for sustaining prolonged hyperemic responses, as well as their heightened susceptibility to inflammatory disruptions. For longer periods of functional hyperemia, systemic inflammation induces relatively greater alterations to wildtype mice compared to AD mice.

### 3.1 Astrocyte Ca^2+^, microvascular blood flow, and AD

After multiple decades, extensive debates continue regarding the precise contributions of astrocytes for regulating resting state cerebral blood flow and functional hyperemia. In general, resting astrocyte Ca^2+^ is understood to help meet basal metabolic needs by precisely maintaining arteriole diameter. During focal increases in neural activity, glutamate-triggered increases of astrocyte cytosolic Ca^2+^ are believed to stimulate release of vaso-active compounds that modulate arteriole and capillary diameters [27,41–45]. Under different experimental conditions, however, astrocyte Ca^2+^ has been shown to induce both dilation and constriction in arterioles [46]. And multiple studies reported previously that Ca^2+^ released from astrocytic soma occurs too slowly to influence arteriole dilation [47,48]. Conversely, other studies emphasize the significance of faster, previously-undetected astrocyte Ca^2+^ transients in astrocyte endfeet and processes for dilation of arterioles, capillaries, and the contractile sphincters connecting these microvessels [29,30,49–52]. Disparate observations are likely attributable to differences in experimental methodology, age, brain region, stimulus paradigm and frequency, and the presence or absence of anesthesia. Our findings in 12-month old awake female mice support more nuanced assertions that, rather than initiating it, astrocytes contribute to modulation and continuation of the stimulus-evoked hemodynamic response, along with its recovery to baseline levels. Although we observed a more pronounced influence of astrocyte endfeet and processes on arterioles compared to higher-order capillaries, the tradeoffs between imaging speed and imaging area precluded more robust measures of faster, low amplitude Ca^2+^ transients [30,50]. In future studies, improvements in imaging acquisition speed could better delineate the presence and influence these faster Ca^2+^ processes, as well as their prospective variations with AD pathology or systemic inflammation.

In previous studies of AD mouse models, both hyperactive and diminished astrocyte Ca^2+^ signaling have been reported. The results vary with brain region, disease stage, the presence or absence of a functional stimulus, and notably, animal model [53–55]. In the absence of induced systemic inflammation, our observations are consistent with prior reports in APP/PS1 mice that observed increased spontaneous release of cortical astrocyte Ca^2+;^, [56,57]. Compared to WT control mice, the magnitude of sensory-evoked astrocyte Ca^2+^ release is larger in AD mice for a brief 3 s whisker stimulus (Fig 1. G-I). However, for a longer 30 s stimulus, the biphasic responses revealed that astrocytes in the AD mouse brain achieve their maximal Ca^2+^ release faster than WT mice but, ostensibly, lack sufficient Ca^2+^ stores to sustain the hyperemic response (Fig. 1K-M). During the latter part of the longer stimulus, we observed that diminished Ca^2+^ release in AD mice correlates with reduced dilation of penetrating arterioles and capillaries and faster return to baseline diameter. Our observations agree well with prior studies in anesthetized 6-month old APP/PS1 mice. Lines et al also observed both increased spontaneous release of Ca^2+^ and decreased evoked astrocyte Ca^2+^ release from 20 seconds of hind-paw stimulation [58]. The precise mechanism(s) of aberrant Ca^2+^ in reactive astrocytes currently remains inadequately understood. Upregulation of the purinergic and the metabotropic receptors, increased mitochondria Ca^2+^ efflux, endoplasmic reticulum Ca^2+^ concentration, and variations in neuronal hyperactivity could all contribute to alterations in spontaneous Ca^2+^ activity within reactive astrocytes [10]. Future studies investigating the internal store concentrations of Ca^2+^ in astrocyte mitochondria and endoplasmic reticula can clarify their influence on functional hyperemia [59].

Impaired neurovascular coupling has been observed in both AD patients and mouse models, including Tg2576, J20, and the triple transgenic 3xTg-AD mouse models [53,60–64]. Our OISI and 2-photon experiments show that APPPS1 mice show mildly reduced stimulus-induced changes to cerebral blood volume and penetrating arteriole diameter (Fig1.D, F, J). Preserved neurovascular coupling was also reported in anesthetized J20 mice modeling familial AD at age 9-12 months [65,66]. In high-order capillaries (diameter < 5 μm), we observed increased resting-state RBC velocity but reduced dilation and faster relaxation time in response to 30 s stimulus (Supplementary Fig. 2 C&H). Gutiérrez-Jiménez et al. found reduced resting-state capillary flux and elevated capillary RBC velocity and arteriole dilation induced by 20 s electrical stimulation evoked in 18-month-old APPswe/PS1dE9 model [67]. Early morphological alterations in microvasculature have also been reported in APPswe/PS1dE9 mice with electron microscopy [68] and in 3x-Tg mice with optical coherence tomography [69]. Taken together, the current and prior observations strongly support emerging hypotheses that vascular impairments in AD originate in the microcirculation before later propagating to larger vessels [70].

### 3.2 Systemic inflammation’s impact on neurovascular coupling and Astrocyte Ca^2+^ signaling

In previous studies with healthy mice and rats, acute periods of LPS-induced systemic inflammation (4-24 hours) yielded substantial reductions in resting state capillary blood flow, cerebral blood volume, and cerebral metabolic rate of oxygen (CMRO_2_) responses during functional hyperemia [23,71,72]. Here, we showed that a prolonged 14-day period of LPS-induced systemic inflammation provoked mild reductions to baseline diameters of precapillary arterioles and capillaries (Supplementary figure 5) and to vascular dilation for brief 3 s whisker stimulation (Fig. 3, supplementary Fig. 3). We observed more pronounced reductions in arteriole dilation in WT and AD mice in response to longer 30 whisker stimulation (Fig. 4, supplementary Fig. 4). Compared to prior reports, our milder observations can be attributed to the lower dose of LPS (0.3 mg/kg, instead of >1 mg/kg). Additionally, the chronic protocol of low-dose LPS administration (< 1 mg/kg) reportedly induces immune adaptations at later phases in C57BL/6J and APP23 mice, indicated by temporal alterations in pro-inflammatory and anti-inflammatory cytokine expression [34,33,35]. The prior findings suggest a diminishing inflammatory response over several days of LPS administration. Our immunofluorescence results support this phenomenon by showing reduced GFAP expression in astrocytes at day 14 (Fig. 5).

Our findings demonstrate that prolonged inflammation diminishes astrocyte Ca^2+^ dynamics more substantially than microvascular hemodynamics. In astrocytes and other constituents of the neuro-glio-vascular unit, acute periods of LPS-induced inflammation have been shown to elicit notable transcriptional changes, alterations to inflammatory cytokine secretion, and cellular excitability [73–77]. To our knowledge, neither acute nor chronic systemic inflammation’s influence on astrocyte Ca^2+^ signaling have been investigated previously. In both AD and WT mice, we observed persistent, compartment-specific reductions in astrocyte Ca^2+^ release under resting state and during functional stimulation. Again, LPS-induced reductions to stimulus-evoked astrocyte Ca^2+^ were more severe for a longer functional stimulus in both AD and WT mice, indicating that a secondary inflammatory threat further compromises the internal store concentrations of Ca^2+^ in astrocytes of AD mice.

Our study compellingly support the hypothesis that AD pathology increases astrocytes’ susceptibility to a secondary inflammatory challenge [73]. Our results highlight the significant, critical link between inflammatory and vascular impairments of preclinical AD pathology and the detrimental “second hit” impact imposed by an acute period of illness. Importantly, Ca^2+^ signaling also reportedly regulates astrocytic inflammatory processes by driving NFAT- and NFκB-mediated transcription. Future studies will focus on the role of astrocyte Ca^2+^ as the bidirectional link between neuroinflammation and cerebral blood flow in AD pathogenesis, as well as its role in augmenting astrocyte phagocytosis of Aβ_1-42_ (Fig. 5) [37,78,36,38,7].

## Supporting information

Experimental Data

## DECLARATION OF INTERESTS

The authors declare no competing interests.

## ACKNOWLEDGMENTS

This work was performed with generous support from the Northeastern College of Engineering and the National Institutes of Health: NIH R01AA27097, NIH R56AG058849, and R21AG085655. We thank Dr. Martin Thunemann and John Jiang from Boston University for their guidance with animal surgery and calcium imaging. We thank Dr. Guoxin Rong and the Institute for Chemical Imaging of Living Systems (RRID:SCR_022681) at Northeastern University for consultation and imaging support, and for the assistance of confocal imaging and wide-field fluorescence imaging. We thank Dana-Farber/Harvard Cancer Center in Boston, MA, for the use of the Specialized Histopathology Core, which provided brain slices immunofluorescence staining service. Dana-Farber/Harvard Cancer Center is supported in part by an NCI Cancer Center Support Grant # NIH 5 P30 CA06516.

## 4. METHODS

### 4.1 Animal preparation

All experiments were performed following ARRIVE guidelines for animal care, under a protocol approved by the Northeastern University Institutional Animal Care and Use Committee. The mouse strain used for this research project, B6C3-Tg(APPswe,PSEN1dE9)85Dbo/Mmjax, RRID:MMRRC_034829-JAX, was obtained from the Mutant Mouse Resource and Research Center (MMRRC) at The Jackson Laboratory, an NIH-funded strain repository, and was donated to the MMRRC by David Borchelt, Ph.D., McKnight Brain Institute, University of Florida. The double transgenic AD mouse co-expresses the Swedish mutation of amyloid precursor protein (APP) and a mutant human presenilin 1 gene (PS1-dE9) and accumulates beta-amyloid deposits in the cortex at 6 months [1]. The study used female APP/PS1dE9 mice and their wild-type littermates (age 12-13 months, n=8 per cohort). At 10 months of age, mice underwent cranial window surgery followed by GFAP-promoted GCaMP6f virus injection. Under isoflurane anesthesia (3–3.5% isoflurane for induction and 1–2% for maintenance), hair and skin was removed to expose the skull, and a titanium butterfly-shaped head post was implanted using dental cement [80]. After the localization of the injection site, a ∼ 3 mm diameter of cranial window was created on the barrel cortex of the mouse, followed by targeted injection of a GFAP-promoted GCaMP6f virus (pZac2.1gfaABC1D-cyto-GCaMP6f, Plasmid #52925, Addgene). Virus (300–500 nL) was injected at 2 nL/s, 300–350 μm below the brain surface, using a glass micropipette and a Nanoinjector III (Drummond Scientific) [81]. Injections were performed in two to three cortical regions to increase the area of viral expression. The procedure lasted 15 to 20 minutes, with the dura kept hydrated throughout. After the injections, the dura was covered with a CNC-machined transparent acrylic coverslip (outer diameter: 5 mm, inner diameter: 3 mm). The mouse was then removed from the stereotaxic frame and allowed to recover from anesthesia. The animal was single-housed and provided with 5 days of post-operative care (40mg/ml sulfamethoxazole (SMX) and 8 mg/ml trimethoprim (TMP) in drinking water). Buprenorphine (0.05 mg/kg at 0.03 mg/ml) was administered subcutaneously for three days after surgery. To label astrocytes in the activation center, we conducted optical intrinsic signal imaging (OISI) and whisker stimulation on the anesthetized mouse before removing the skull. The skull over the barrel cortex (A-P: 2mm, M-L: 3 mm) was thinned, and whiskers on the contralateral side of the thinned skull were deflected using a cotton swab. The skull was illuminated with an LED light source during the stimulation through a green bandpass filter (568/10 nm). The reflected light was recorded with a CMOS camera (Basler acA1300– 200*µ*m, Edmund optics) mounted on the surgical microscope, and the relative intensity change was calculated with a customized MATLAB script. Regions with the greatest intensity drop were identified as the activation area and selected as the site for viral injection.

### 4.2 Habituation training

Headpost restraint training started 7 to 10 days post-surgery using a custom-made imaging cradle. During training, the mouse was conditioned to tolerate head immobilization from 5 minutes to 1 hour. Whisker stimulation was included during the final training sessions to acclimate the animal to the functional activation experiment. Milk was given as a reward during the training.

### 4.3 Systemic inflammation induction

Peripheral inflammation induced by systemic injection of LPS has been shown to transfer to the central nervous system and cause a prolonged inflammatory response in the brain [82]. Systemic inflammation was induced through daily intraperitoneal (i.p.) injections of 0.3 mg/kg lipopolysaccharide (LPS, *E. coli* O55: B5, L4005, CAS-No. 93572-42-0, Sigma-Aldrich) for 14 days. A stock solution of LPS was prepared at 1mg/mL in phosphate-buffered saline (PBS) and stored at −20°C. The solution was vortexed and diluted 4-fold to 0.25 mg/mL for each injection.

### 4.4 Functional activation protocol

A capillary tube with a 3D-printed broom-like end was positioned approximately 2.5–3 cm from the nose, parallel to the right side of the mouse’s face, and below the eye to avoid directing air toward the eyes during stimulation. Air puffs were delivered through the tube and controlled using a pneumatic drug injection system (PDES-DXH, ALA Scientific, USA). Two whisker stimulation protocols—brief and prolonged—were performed. The brief protocol consisted of a 5-second baseline, 2.7 seconds of 1.67 Hz stimulation (5 pulses, 20 psi), and 22.3 seconds of recovery. The prolonged protocol included a 5-second baseline, 30 seconds of 1.67 Hz stimulation (50 pulses, 20 psi), and 30 seconds of recovery. Short and long stimulations were interleaved within each trial and repeated 10 times.

### 4.5 Functional activation center mapping

The activation center of whisker stimulation was mapped using optical intrinsic signal imaging (OISI) under the brief whisker stimulation protocol. A white LED illuminator (SCHOTT KL 1600) with a 568/10 nm bandpass filter (Edmund Optics) served as the light source. The camera was triggered by a TTL signal from a general-purpose input/output (GPIO) hardware trigger. A fiber-optic LED light source illuminator (SCHOTT KL 1600 LED) delivered light to the cranial window at a 60° angle. Reflected light was collected using a 4× objective (Nikon Instruments Inc.) and recorded at 5 Hz for 20 seconds with a CMOS camera (Basler acA1300–200μm, Edmund Optics). Baseline intensity (I□) was calculated by averaging the intensity over the baseline recording. Relative intensity (I/I□) was calculated for all imaging frames. Cortical area with the maximal intensity decreases was identified as the activation center.

### 4.6 Two-photon imaging in awake mice

Two-photon imaging of astrocyte Ca^2+^ signaling and vascular dynamics at barrel cortex at resting-state and during functional activation was conducted using the multiphoton imaging system Ultima2pPlus (Bruker Nano, Inc) with a 25× objective (1.10 NA, Nikon Instruments Inc). Imaging during the functional activation experiment was synchronized with the whisker stimulation protocol. Blood vessels were labeled through retro-orbital injection of Texas Red Dextran (2.5% w/v, 80uL). A tunable ultrafast laser (InSight X3+, Spectral-Physics) was tuned to 940 nm for simultaneous imaging of blood vessels and astrocytic Ca^2+^ imaging. We tracked the penetrating arterioles from the pial surface to the deepest assessable cortical layer and recorded the vessel diameter and the astrocyte Ca^2+^ release using a minimal frame rate of 4.3 Hz. The image field of view (FOV) was around 88 × 88 *um*^2^. On the following day, capillary diameter and red blood cell velocity in response to the interleaved functional activation was measured with two-photon line scanning [83], with a scan frequency ranging from 369.87 to 1034.8 Hz. Penetrating arterioles perpendicular to the image plane and capillaries parallel to it were selected to ensure accurate diameter estimation, while tilted vessels were excluded during post-processing. We measured regions from 49 to 414 *µ*m below the pial surface for WT mice and 30 to 360 *µ*m below the pial surface for AD mice.

### 4.7 Two-photon imaging data analysis

*Astrocyte spontaneous Ca^*2+*^:* Image frames were pre-processed using a non-rigid motion correction algorithm [84], followed by median filtering. Resting-state astrocyte Ca^2+^ fluctuations were quantified using a ROI-based Ca^2+^ analysis tool: “GECIquant” in ImageJ [85]. Briefly, average intensity projection was generated from the time series of two-photon raster scan images. We created a binary region of interest (ROI) mask where the area within the polygon ROI was set to 1 and the pixels in the rest of the image was set to 0. Mean intensity within the ROI over time was calculated. The area criterion was set to 30 *µm*^2^ to Inf for astrocyte soma and endfeet, and 8 to 2000 *µm*^2^ for Ca^2+^ waves in processes. Relative intensity change (Δ*F*/*F*_0_ = (*F*_*t*_ - *F*_0_)/*F*_0_) was calculated. *F*_0_ was calculated as the median of the lowest 5% fluorescence intensity over the entire 10-minute raster scan. Δ*F*/*F*_0_ of all astrocyte ROIs from all mice were interpolated with a d*t* = 0.2244 *s*, (i.e., 4.45 fps). The Δ*F*/*F*_0_ traces were pre-processed using linear detrending to remove intensity drift that occurred during the measurement. The detrended intensity traces were then low pass filtered with a cutoff frequency of 0.1 Hz, as determined by power spectrum analysis. To determine the threshold for the Ca^2+^ event, we sorted the normalized intensity values in ascending order, calculated the median absolute deviation (MAD) of the lowest 80% of the Δ*F*/*F* values, and used 5 times the MAD as the threshold for the Ca^2+^ event. This threshold is robust to outliers and helps to reduce the number of ‘false positives’. Ca^2+^ events were detected and quantified using the ‘findpeaks’ function in MATLAB. The detected events were categorized as either ‘singlepeak’ or ‘multipeak’ events as described [86]. We calculated the frequency of the event as per min per ROI.

*Arteriole diameter:* Arteriole diameter in response to functional activation was quantified using the automatic structure tracking algorithm developed by Haidey et al. [87]. The algorithm tracks the cross-sectional area of the vessel based on user-defined luminal edges. Vessel diameter (D) was calculated from the cross-sectional area. Relative change of diameter (Δ*D*/*D*_0_ = (*D*_*t*_ - *D*_0_)/*D*_0_ × 100) was calculated, where the baseline diameter (*D*_0_) was the average diameter over the 5s resting-state measurement. Since the frame rates of different imaging fields sometimes vary slightly, Δ*D*/*D*_0_ were interpolated with Δ*t* = 0.2244 *s*.

*Quantifying stimulus-evoked responses:* To quantify stimulus-evoked arteriole dilation and astrocyte Ca^2+^ release, we first scrutinized all 10 stimulation trials and excluded those with motion artifacts. The remaining trials were classified as either ‘with response’ or ‘without response.’ Trials exhibiting responses were averaged to determine the mean ΔD/D0 (arteriole dilation) and ΔF/F0 (Ca^2+^ activity) for each region of interest (ROI). The averaged arteriole dilation and Ca^2+^ signals were low-pass filtered with a cutoff frequency of 0.2 Hz. Peak responses were identified using the ‘findpeaks’ function in MATLAB. For each detected peak, the amplitude and latency were calculated. Peak latency was defined as the time interval between stimulus onset and the peak response. The area under the curve (AUC) was calculated for each stimulation paradigm, as reported by previous publication [28]. For 3s whisker stimulation, AUC was computed over the 0–25 s window, while for 30s stimulation, it was assessed over 0–60 s. Additionally, for 30s stimulation, the average response during the last 2s of stimulation was calculated to assess late phase activity. To analyze response kinetics, we followed the method used in a recent publication to fit ΔD/D □ and ΔF/F □ traces with a sigmoidal model [88]. The rise part of the trace was fitted using the equation 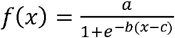 and the decay part of the trace was fitted using the 4-parameter logistic equation 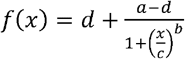 Rise time was calculated as the interval between 10% and 90% of the fitted peak response amplitude. Fall time was determined as the time required for the signal to decay from 90% to 10% of the peak response in the fitted curve. Duration was calculated as the full width half maximum (FWHM). Note that responses to 30s stimulation have a plateau phase, the decay part of the traces was defined when the responsive trace start to return to the baseline. Duration for 30s stimulation is determined as the time interval from the half-maximal of the rising phase to the half-maximal of the decay phase.

*Capillary diameter:* Space-time images from two-photon line scans were denoised using a 2D median filter. Vessel diameter was calculated using the MATLAB two-photon imaging analysis toolbox, CHIPS [89]. The algorithm normalizes the vessel’s intensity profile to a range of 0 to 1 and calculates the diameter using the full width at a user-defined height. We used the full width at 0.4 of the maximum intensity, as it provided the most robust diameter estimation based on our signal-to-background ratio. The sampling rate was set at a maximum of 20 Hz. Measurements affected by motion, particularly along the axial axis were discarded. Average diameter and ΔD/D□ across all selected stimulation trials were calculated. The ΔD/D□ traces were smoothed using a moving median filter with a one-second window (20 data points). Vascular dilation amplitude, AUC and kinetics were analyzed using the same method as in raster-scan data analysis. Vessels were categorized by branch order: penetrating arterioles (0^th^-order), pre-capillary arterioles (1^st^ and 2^nd^order), and higher-order capillaries (>3^rd^-order branches). Resting-state capillary diameter was calculated by averaging the baseline measurement taken from 1.5 s to 4.5s.

*Capillary RBC velocity:* The space-time images were filtered using Sobel filtering, and the capillary red blood cell (RBC) velocity was analyzed using the iterative Radon Transform [90]. Image segments of every 100 lines were used to calculate RBC velocity. 25 lines were skipped before calculating the velocity at the next time point, leading to a time window of the RBC velocity from 24 ms to 67 ms. Relative velocity (Δv/v□,%) was calculated and filtered with a low-pass filter with a cut off frequency of 0.1 Hz. All Δv/v□ traces were interpolated with dt = 0.05s.

*Line scan measurement of astrocyte Ca^2^:* Line segment measuring the astrocyte processes and endfoot wrapping around the capillaries were selected during post-processing. Total intensity along the line was calculated and normalized to ΔF/F0 (%). The normalized intensity traces were also downsampled to 20Hz. Amplitude, AUC and kinetics were quantified.

### 4.8 Ex vivo immunofluorescence imaging (IF)

After 14 days of LPS treatment, mice were sacrificed, and brains were collected for immunofluorescence (IF) staining. IF staining was performed on a subset of the imaged mice (n=3 per cohort). Mice received 14 days of saline injections at a volume equivalent to the LPS dose of 0.3 mg/kg served as control group (n=2 for WT, n=3 for AD). A separate group of mice received 7 days of LPS injection was used as well (n=2 for AD, n=3 for WT). Mice were intracardially perfused with 40 mL of heparin-PBS solution, followed by 20 mL of 4% paraformaldehyde (PFA) for fixation. The brains were collected and stored in 4% PFA for a maximum of 24 hours, then transferred to 70% ethanol and stored in −4 ° C fridge. The brain samples were sent to the Dana-Farber/Harvard Cancer Center Specialized Histopathology Services Core (Boston, MA, USA) for immunofluorescence staining. The samples were paraffin-embedded and sliced into approximately 10 μm thick sections for IF staining. Nuclei were stained with DAPI, microglia with an anti-Iba-1 antibody labeled with Alexa Fluor 594, and astrocytes with an anti-GFAP antibody labeled with Alexa Fluor 647. IF images were collected with confocal microscopy (LSM 800, ZEISS). DAPI, AF-647, and AF-594 were excited with 405nm, 640nm, and 561nm, respectively. Images were taken using 5× and 10× objectives. Barrel cortex and hippocampus were identified using the Allen Brain Atlas. For each 10× field of view (638 × 638 *um*^2^), Z-stack images were collected with a step size of 0.34 *µm*. All images were acquired using consistent excitation power and emission detector gain settings. Aβ_1-42_ was stained with anti-Aβ_1-42_ antibody conjugated to Alexa Fluor 647. Fluorescence images of Aβ_1-42_ were acquired with wide-field fluorescence microscopy (Zeiss Axio Observer). All images were acquired using the same exposure time.

GFAP and Iba-1 expression were quantified using the maximum intensity projection (MIP) of the z-series images. Background was removed and MIP images were binarized using an intensity threshold. The percentage area of pixels with intensity above the intensity threshold was calculated. Similarly, percentage area of Aβ_1-42_ load in cortex and hippocampus was quantified in Image J.

### 4.9 Statistical analysis

We performed statistical tests to see if there is difference in spontaneous astrocyte Ca^2+^ dynamics, stimulus-evoked vessel dilation and astrocyte Ca^2+^ release in AD, and how LPS-induced systemic inflammation affects these parameters in both WT and AD cohorts. Differences between WT and AD group before LPS injection were tested using the Wilcoxon rank sum test on MATLAB. Effect of LPS on both cohorts were tested using linear mixed effect model with the ‘lme4’ package in RStudio. Time (i.e., day0, day 7 and day14) was set as the fixed effect. Pair-wise comparison was conducted with Tukey’s method in the ‘emmeans’ package. * = p<0.05, ** = p<0.01, ***=p<0.001, ****=p<0.0001.

## Notes

### Competing Interest Statement

The authors have declared no competing interest.

### Summary of Updates

Supplemental files updated. Added experimental data sets.

